# Portuguese wild grapevine genome re-sequencing (*Vitis vinifera sylvestris*)

**DOI:** 10.1101/2020.02.19.955781

**Authors:** Miguel J N Ramos, João L Coito, David F Silva, Jorge Cunha, M Manuela R Costa, Sara Amâncio, Margarida Rocheta

## Abstract

*Vitis* is a relevant genus worldwide. The genome of a *Vitis vinifera* representative (PN40024) published in 2007 boosted grapevine related studies. While this reference genome is a suitable tool for the overall studies in the field, it lacks the ability to unveil changes accumulated during *V. vinifera* domestication. Considering that grapevines for wine production (*V. v. vinifera*, hereafter *vinifera*) have evolved from *V. v. sylvestris* (hereafter *sylvestris*), or from a shared no-longer existing ancestor, both subspecies are quite close, but *sylvestris* has not been domesticated and still exist nowadays, preserving wild characteristics, making it a good material to provide insights into *vinifera* domestication. The difference in the reproductive strategy between both subspecies is one of the characteristics that sets them apart. While *vinifera* flowers are hermaphrodite with functional male and female organs, *sylvestris* is mostly dioecious. Male plants present flowers lacking functional carpels unable to produce grapes and female individuals have flowers with reflexed stamens producing infertile pollen but able to exhibit small and acidic grapes. In this paper, we describe the re-sequencing of the genomes from a male and a female individual of the wild *sylvestris* and its comparison against the reference *vinifera* genome.

## Introduction

*Vitis vinifera vinifera* (hereafter *vinifera*) is a worldwide important crop, mainly due to wine production. *Vinifera* has already two sequenced genome versions^1,2^, obtained from PN40024 genotype, a Pinot Noir^1^ variety back crossed to reach 93% homozygosity. The contribution of these *vinifera* genomes knowledge is unquestionable, however, they are limited in providing information on particular changes that occurred during domestication. One of the most relevant subspecies of *V. vinifera* history is *Vitis vinifera sylvestris* (hereafter *sylvestris*), which some authors have referred as *vinifera*^3,4^. *Sylvestris*, free from human selection, may provide clues to explain *vinifera* domestication. This human intervention was an economically important step for grapevine, responsible for morphological changes that include larger berry and bunch size, higher sugar content, altered seed morphology, and a shift from dioecy to a hermaphroditic mating system^5^. The dioecious *sylvestris*, produce female individuals with reflexed stamens, where infertile pollen is produced, whereas male individuals are unable to produce grapes as flowers lacks a functional carpel^6^.

In the *Vitis* genus, a number of studies have contributed to partially uncover the genomic regions that may be responsible for the differences in flower development between hermaphrodites and dioecious plants^7–9^. However, the identified regions (chr2:4,907,434..5,050,616)^7,8^ are poorly sequenced, assembled and annotated in the canonical reference genome^1–3,5,9^. Recently, two new studies^10,11^ suggested the possible players involved in *sylvestris* sex specification by uncovering *INAPERTURATE POLLEN1* (*INP1*) as a male sterility gene and proposing a set of genes responsible for female sterility.

The aims of this paper are: (1) to describe the re-sequencing of *sylvestris*, considering two distinct individuals, one with male flowers and a second one with female flowers; (2) To compare in detail the genomes of the *sylvestris* individuals to the reference genome, in terms of Single Nucleotide Polymorphisms (SNPs) and Insertions/Deletions (InDels); (3) To compare the different alleles found for each *sylvestris* specimen, focusing on chromosome 2, where the sex specification region has been proposed to be present.

## Results

The genomes of two wild-type individuals (*sylvestris*) were obtained by Illumina (2×125) and the reads were mapped against the reference genome of *vinifera* PN40024^**1,2**^ (Figure 1). The reference genome was firstly published in 2007 after controlled breeding of Pinot Noir, creating a lineage with about 93% homozygosity^**1**^. This genome (8X coverage) consists in 487 Mb spread into 3,514 scaffolds (N_50_ = 2.07 Mb)^**1**^. Scaffolds were further associated within the 19 chromosomes of *Vitis* plus 13 virtual chromosomes, corresponding to 12 random chromosomes (associated to the real chromosomes 1, 3, 4, 5, 7, 9, 11, 12, 13, 16, 17 and 18) and a completely unknown chromosome, where sequences that were not fully associated with a real chromosome were deposited^**1**^. The sequenced genome was further improved with a 12X coverage and subsequent gene annotations were published^**1,2**^.

**Figure 1.**
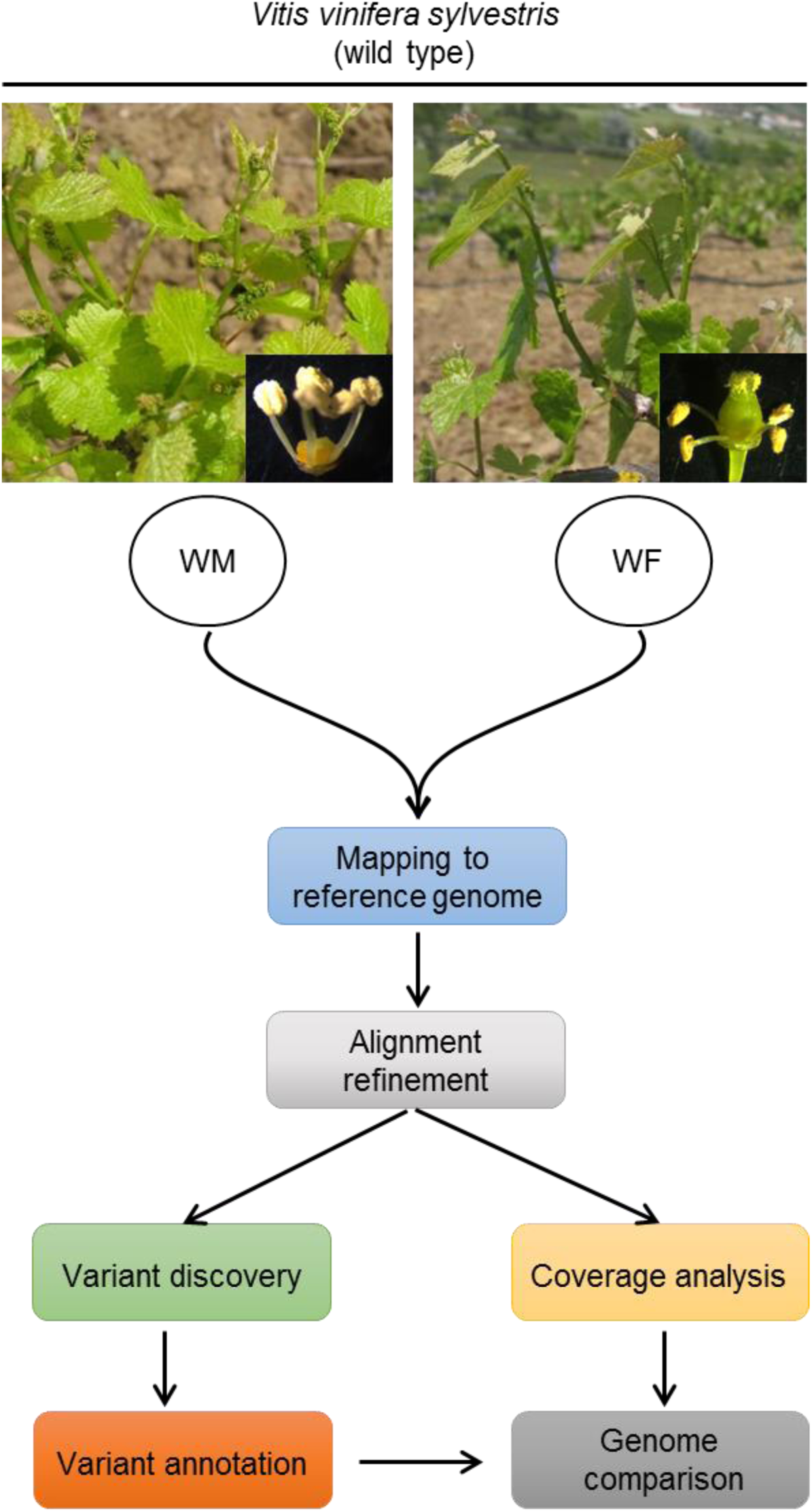
Sampling material of *Vitis vinifera sylvestris* (wild type) and flowchart of data analysis steps. WM individual show carpel abortion (typical of a male flower type); WF individual has reflexed stamens (representing a female flower type). Reference genome is PN40024 (CRIBI, 12X).

### The majority of the wild-type reads map to the reference genome

Around 97% of the high quality reads obtained were successfully mapped to the reference genome: 478,309,356 reads in total (Table 1). From those, 89% mapped only once to the reference genome (Table 1). About 10% of the mapped reads lost their pair in the process or mapped onto a different chromosome (Table 1).

**Table 1.** Number of wild-type reads mapped against the reference genome (PN40024). Unique: reads that map only to one site on the reference genome. Non-Unique: reads that map to multiple sites of the reference genome. Singletons: reads that map to the reference genome, without mate pair. Cross-contigs: reads whose pair map to a different contig. Percentages: percentage of each class to the total mapped reads.WF: Reads obtained from the female wild individual. WM: Reads obtained from the wild type individual with male flower types.

| **Read category** | **WF** | **WM** | **Total** |
| --- | --- | --- | --- |
| Total mapped reads | 209,622,690 | 268,686,666 | 478,309,356 |
| Unique | 187,424,438<br>(89.41%) | 240,250,914<br>(89.42%) | 427,675,352<br>(89.41%) |
| Non-Unique | 22,198,252<br>(10.59%) | 28,435,752<br>(10.58%) | 50,634,004<br>(10.59%) |
| Singletons | 1,782,319<br>(0.85%) | 2,151,281<br>(0.80%) | 3,933,600<br>(0.82%) |
| Cross-contigs | 20,897,087<br>(9.97%) | 24,790,708<br>(9.23%) | 45,687,795<br>(9.55%) |

### Evaluation of the differences between the wild-type and reference genomes

Single Nucleotide Polymorphism (SNP) and Insertions/Deletions (InDels) analysis were performed between the sequences obtained for the two wild-type individuals and the reference genome. Wild-type individuals used on this re-sequencing project present distinct flower characteristics. One of the plants show male characteristics (WM), with no functional carpel and the other exhibits female flowers (WF) with a complete carpel, but with reflexed stamens producing infertile pollen. Both of these individuals had almost the same number of SNPs identified (6,197,145 for WF and 6,575,026 for WM individual), and shared half of the occurrences (3,643,713 SNPs; Figure 2a). The InDels analysis showed the same tendency as the previous results: the number of occurrences in both individuals were similar (1,151,997 for WF and 1,177,918 events for WM) and about half of them were shared (637,320 events; Figure 2b).

**Figure 2.**
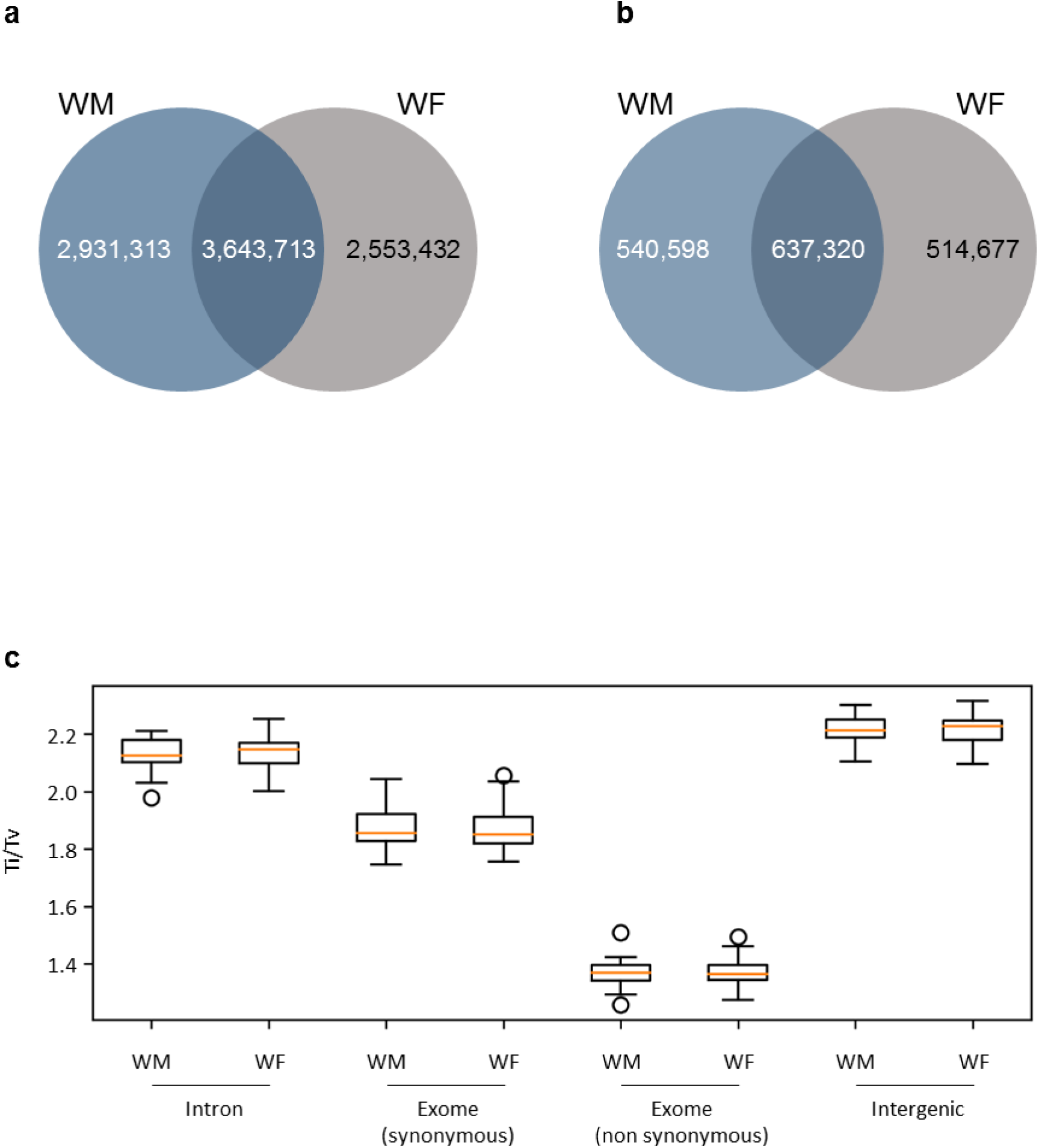
Landscape of SNPs and InDels in the two wild-type genomes. (a) Venn diagrams showing the number of SNPs. (b) Venn diagrams showing the number of InDels. Comparison was performed against the reference genome (PN40024, CRIBI, 12X). (c) Transition over transversion ratio (Ti/Tv) of WM (male) and WF (female) individuals according to the genomic region. For exomes, analysis was split into synonymous and non-synonymous SNPs.

The ratio between the numbers of transitions over transversions (Ti/Tv) is usually calculated in order to assess the quality of an assembly or SNPs calling. As Ti/Tv ration has been reported dependent of the genomic context of the SNPs^12^, we have evaluated Ti/Tv on introns (ratio 2.1), exons with synonymous SNP (1.9), exons with non-synonymous SNPs (1.4) and intergenic regions (2.2) (Figure 2c). Our analysis confirmed that Ti/Tv was dependent of the genomic location, but not on the individuals on study, as WM and WF individuals presented the same Ti/Tv pattern (Figure 2c).

We have pooled both sequenced individuals together and considered them as biological replicates in subsequent analysis. The distribution of SNPs and InDels, on the genome, was evaluated for each chromosome, by calculating the frequency of occurrences (number of SNPs/InDels by chromosome size). In both SNPs and InDels, the chromosome 6 was the one with the lowest number of events. For this chromosome, 1.00 % of the bases were found to be changes on a single nucleotide (frequency of 0.011; Figure 3a) and InDels were identified in 0.20% of the chromosome (frequency of 0.0020; Figure 3b). The least conserved chromosome was the 9 with a percentage of SNPs of 1.5% (frequency 0.015; Figure 3a) and InDels covering 0.29% of this chromosome size (0.0029; Figure 3b).

**Figure 3.**
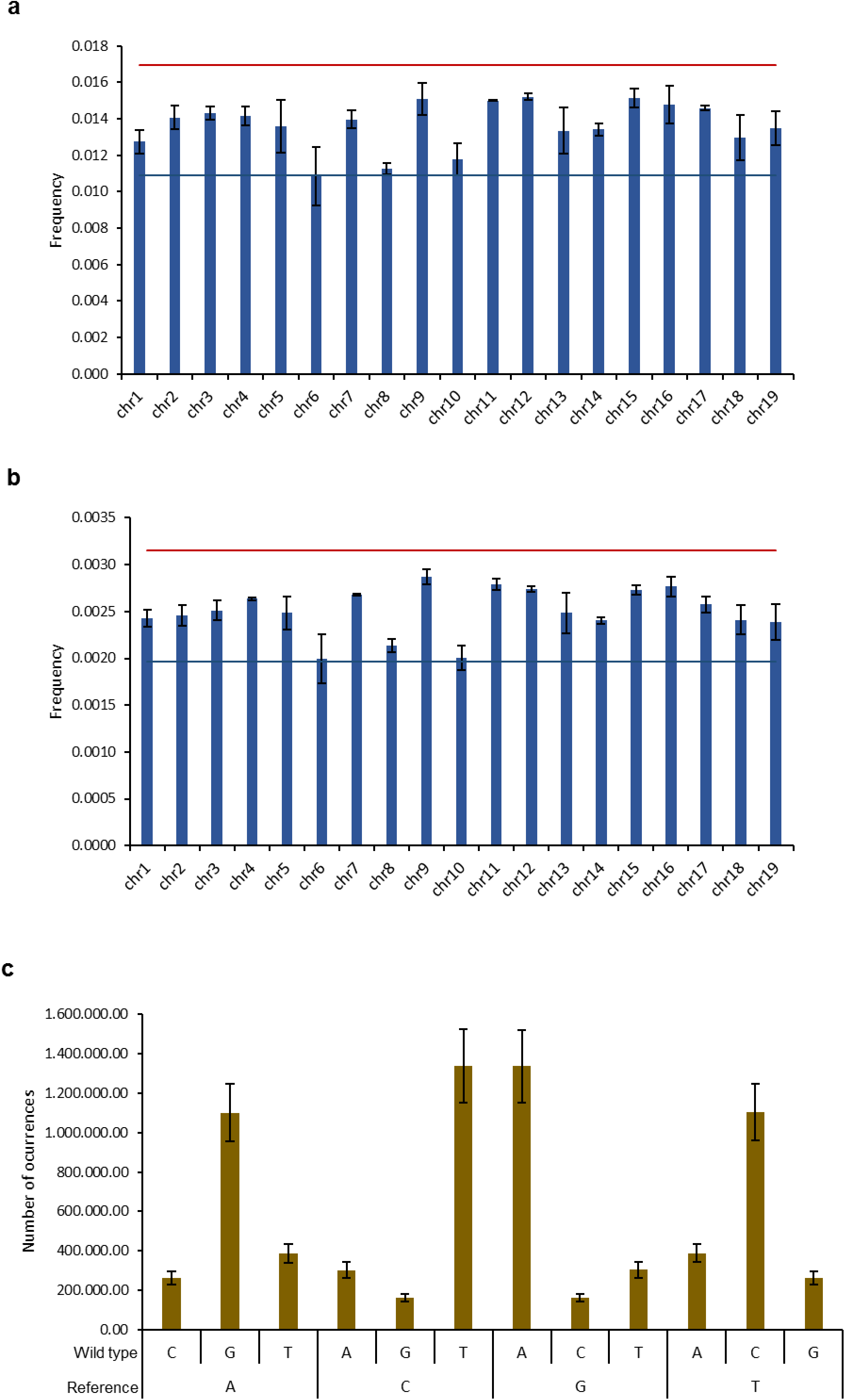
Analysis SNPs distribution and dynamic. (a) Frequency of events per chromosome length (base pairs) in considering SNPs and (b) InDels. Red and blue lines represent the Tukey’s outlier fences (k = 1.5). Vertical bars represent the standard error. (c) Number of transversions (A ↔ C, A ↔ T, C ↔ G and G ↔ T) and transitions (A ↔ G and C ↔ T). Wild type: the nucleotides identified in the wild-type individuals; Reference: The nucleotide in the reference genome (PN40024, CRIBI, 12X);

We analysed the nature of the substitutions in SNPs by comparing the nucleotides on the reference genome with the nucleotides observed for the wild type. In accordance with other studies^10,12^, the number of transitions (substitution between adenosine and guanine or cytosine and thymine) were much more frequent on both genomes than transversions (substitutions between adenosine and cytosine, adenosine and thymine, cytosine and guanine or guanine and thymine) (Figure 3c).

The effects of SNPs on protein-coding regions were also investigated by analysing the SNP location within gene context and the impact caused. The large majority of the SNPs occur in non-coding regions, as intergenic, upstream, downstream and introns (2,240,613;1,891,608; 1,048,262 and 749,096 respectively; Figure 4a). Less frequently, SNPs were identified in exons and UTR regions. Interestingly, the number of SNPs located in exons, with synonymous (104,970) and non-synonymous (138,468) impact was almost the same (Figure 4a). From the impacts that a SNP may cause on protein sequence, premature start and stop codons as well as the loss of start or stop codons (related with the start and stop position in the reference annotation) are among the most disruptive ones. *Sylvestris* genomes had 3,553 premature start codons and 3,183 premature stop codons (Figure 4a), compared with the reference genome. In terms of codons loss, we have identified 530 loss of start codon and 705 loss of stop codon (Figure 4a).

**Figure 4.**
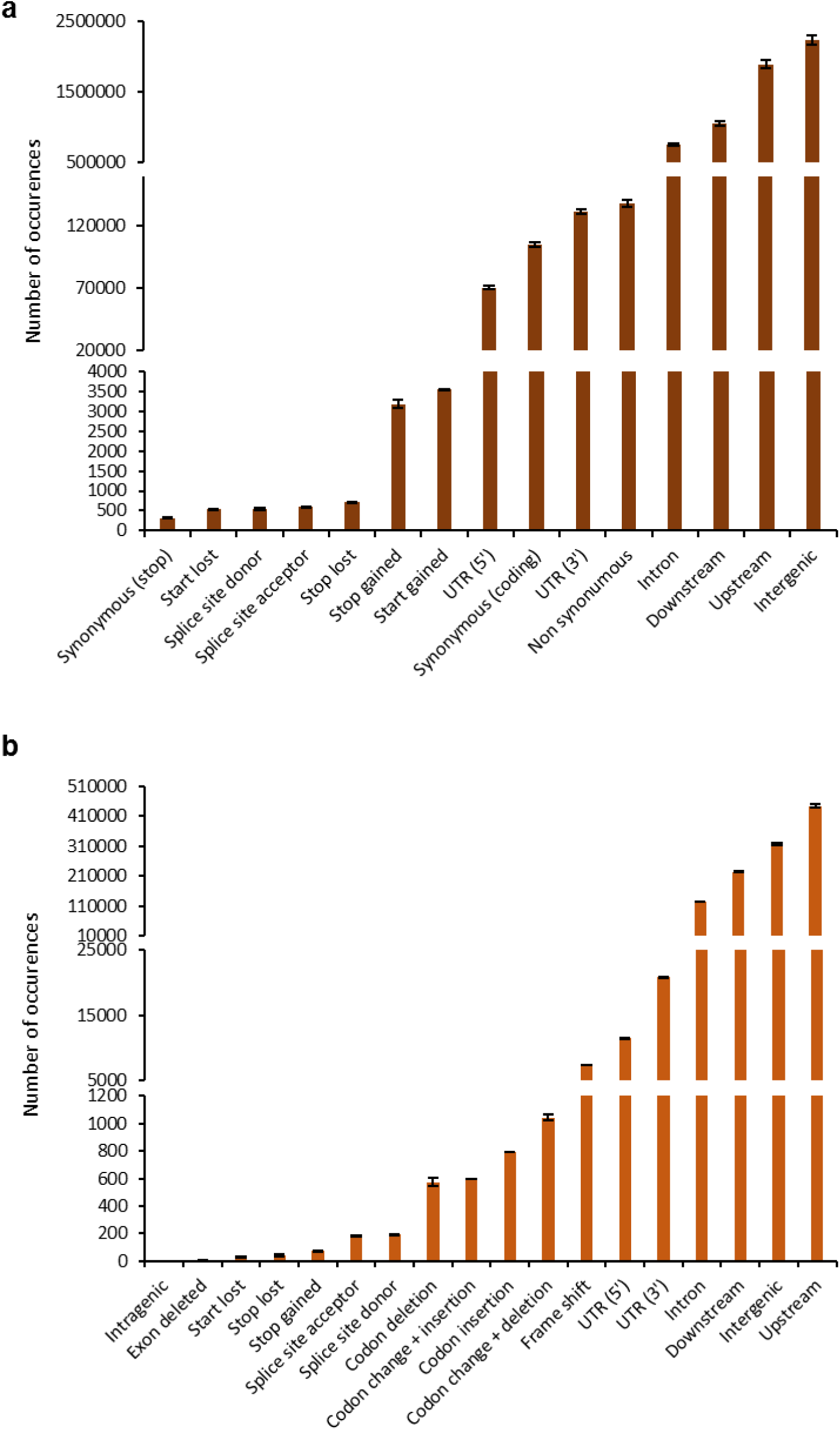
The gene context of the identified modification in wild-type genomes. (a) SNPs. (b) InDels. Reference genome and annotation: PN40024, CRIBI, 12X V2.1. When context is an exon, the impact on protein sequence was evaluated. For events between genes, the most relevant context was chosen (e.g. Upstream instead of Downstream), as described by SnpEff documentation.

The majority of InDels also occurred on non-coding regions (upstream [444,281], downstream [224,906]) (Figure 4b). The few InDels that occurred inside genes are not in coding regions (intron [126,143], 3’-UTR: [20,852], 5’-UTR [11,490]) (Figure 4b). InDels resulting in frame shifts (7,481), codon change + deletions, corresponding to events where at least one codon is changed and at least one is deleted (1,044), codon insertion (792), codon change with insertions (600) and simple codon deletion (573) were also found (Figure 4b). Events as splice site acceptor, splice site donor, start lost, stop lost and stop gained were much less frequent (less than 200 occurrences per event) (Figure 4b). There were also four exon losses in the wild-type individuals studied (Figure 4b)

### Heterozygosity levels

The nature of the SNPs and InDels was evaluated regarding their homozygosity. The majority of the events were heterozygous (0/1 and 1/2) (Figure 5). The *sylvestris* individuals had about 4,104,719 heterozygous SNPs and 761,638 heterozygous InDels. The number of homozygous alterations between *sylvestris* and the reference genome (1/1) was much lower (2,281,366 homozygous SNPs and 403,321 homozygous InDels) (Figure 5).

**Figure 5.**
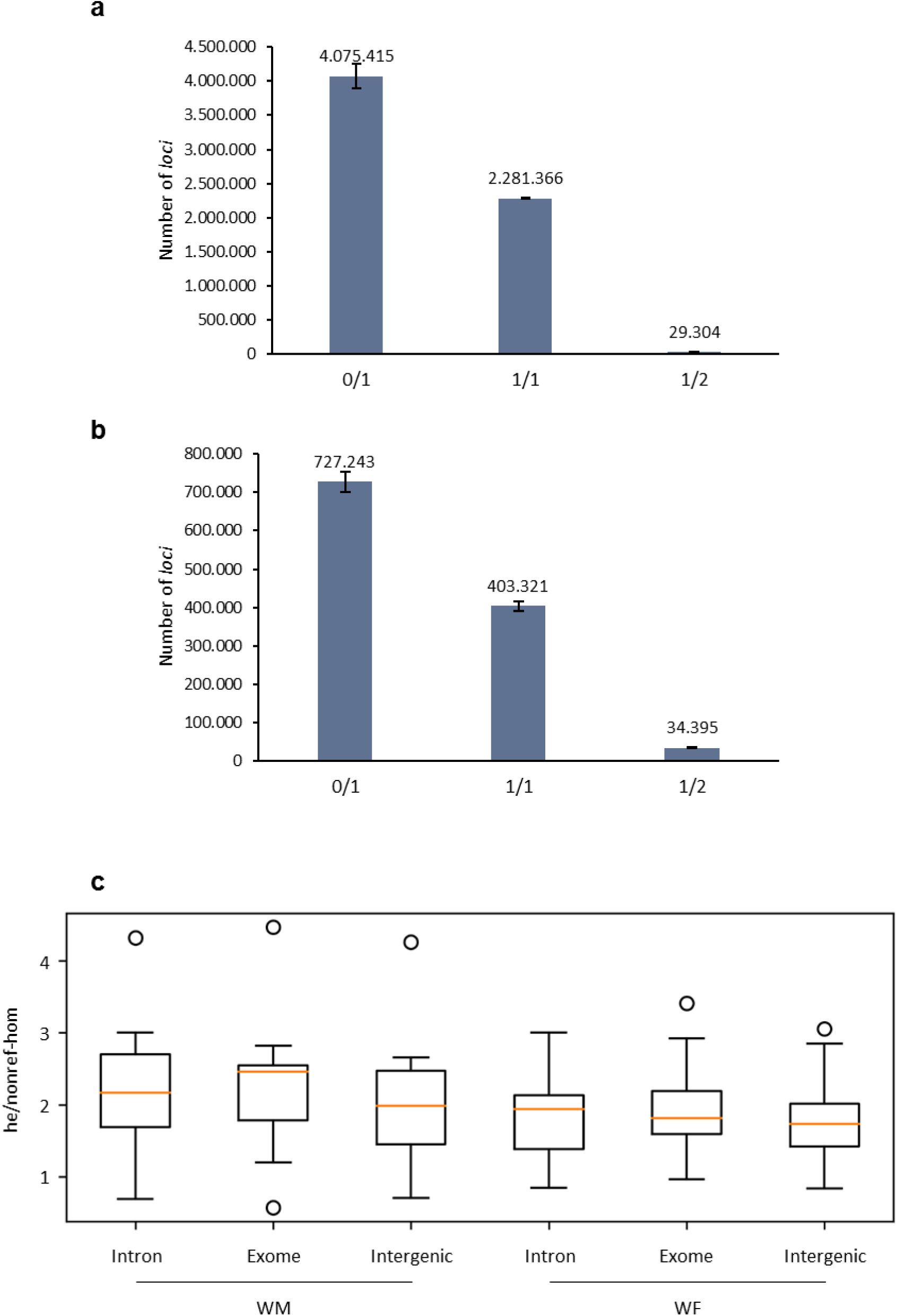
General heterozygosity analysis. Number of homozygous (1/1) and heterozygous (0/1 and 1/2) *loci* evaluated for the wild type re-sequenced genomes. (a) Evaluation according to SNPs. (b) Evaluation according to InDels. (c) Heterozygous over non-reference homozygous ratio (he/nonref-hom) for each individual (WM and WF) according to multiple genomic locations. 0/1 represents *loci* where two alleles were observed, one being identical to reference genome (PN40024, CRIBI, 12X); 1/1 represent *loci* where only one allele was observed, being different than reference; 1/2 represents heterozygous *loci*, being both alleles different than reference.

Combining SNPs and InDels, the total number of heterozygous *loci* (0/1 and 1/2) was 4,866,357 while the homozygous (1/1) changes were 631,487. The ratio between heterozygous/homozygous SNPs was 1.80 and1.89 for InDels.

An analysis of the heterozygous (0/1 and 1/2) over non-reference homozygous (1/1) SNPs, considering the *sylvestris* individuals separated, at multiple genomic locations, revealed slightly differences between them (Figure 5c). Ratios obtained were 2.2 for WM (introns and exons), 2.0 for WM (intergenics), 2.0 for WF (exome) and 1.8 for WF (introns and intergenics) (Figure 5c). No differences were obtained between synonymous and non-synonymous exons (data not shown).

### Validation of genome sequencing by Sanger analysis

Primers were designed to amplify regions where differences have been detected between wild-type sequences and the PN40024 reference^1^. From the nine modifications tested, four were homozygous and shared by both individuals, suggesting that these modifications are exclusive of wild-type individuals (Supplementary Figure S1a, b and c). One of the modifications shared by both individuals represented a case where the wild type genomes are heterozygous (Supplementary Figure S1d). We also validated the occurrences of individual exclusive modifications, where one of the individuals (WF) was homozygous and the other (WM) was found to be heterozygous (Supplementary Figure S1d).

### Manually inspection of chromosome 2

Chromosome 2 has been associated to *Vitis* sex specification^7–9^. The 24 genes located in the region associated with sex were manually inspected from a *de novo* assembly performed with the raw data obtained for both male and female *sylvestris* individuals. We also manually inspected several genes on chromosome 2, some of them described to be involved in flower development like *NGA1* (VIT_202s0025g03000)^15^ and *SLK2* (VIT_202s0025g04390)^16^. Other 20 genes located on chromosome 2, with distinct transcription patterns were previously^15^ detected^16^ and were subject to this analysis, in a total of 46 genes. This analysis was possible since both individuals have a distinct flower phenotype. From the 46 genes analysed, the male individual studied was heterozygous for 25 genes (Figure 6). The alleles of three of these genes were identical to the reference genome (VIT_202s0241g00140, VIT_202s0154g00230, VIT_202s0033g00020; Figure 6). In the female individual, 8 out of 46 genes were heterozygous with both alleles presenting differences compared to the reference genome; only one of 46 genes was identical to the reference genome (VIT_202s0025g02630); and the remaining 37 genes were homozygous with differences when compared to the reference. Also, we found only one identical gene between male and female individuals (VIT_202s0154g00030) being homozygous, but distinct to the reference genome (Figure 6).

**Figure 6.**
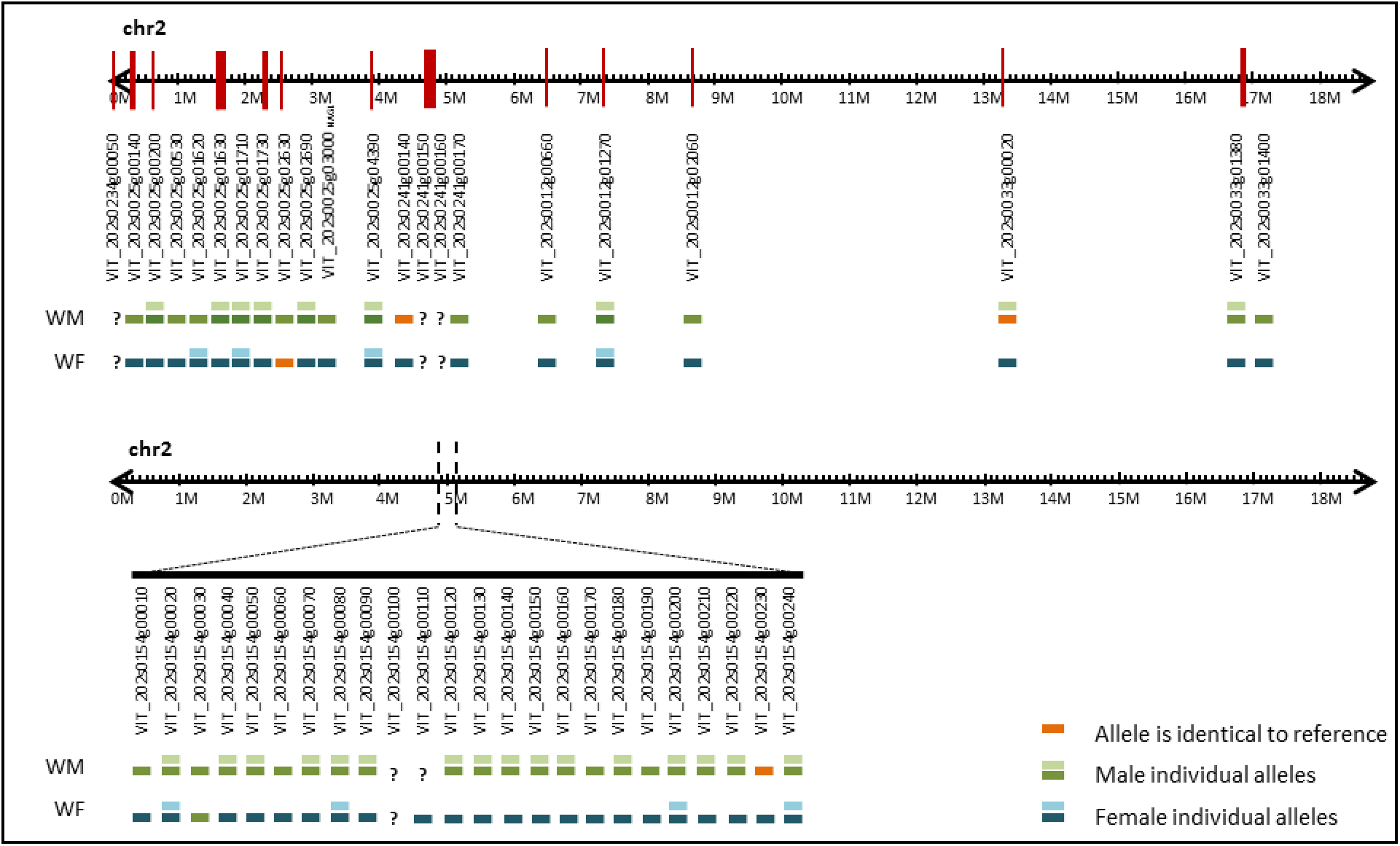
Representation of location and differences between the most relevant genes and sex-associated *loci* in chromosome 2 after manual validation. Orange bars represent alleles of the *de novo* assembled genomes identical to the reference (PN40024, CRIBI, 12X). Green bars represent alleles identified in the male individual. Blue bars indicate distinct alleles observed in the female individual. When a male and a female allele were identical, the same colour was used (green). A single bar was used to represent homozygous genes. Question marks refer to genes with low assembly resolution whose sequence was not identified correctly.

While preparing this manuscript, two independent studies were published regarding the sexual specification of *sylvestris*^10,11^. Both studies indicate that the gene coding for INAPERTURATE POLLEN1 (INP1) is responsible for male sterility, due to an 8-bp deletion in the CDS region, detected in female individuals, causing a premature stop codon. We found that this gene has homology with VIT_200s0233g00051 and VIT_202s0154g00120, associated with chromosomes “unknown” and 2, respectively. We consider that these two IDs are two alleles of the same gene. In the samples tested under this re-sequencing project, we identified the two alleles in the male individual, but the female individual only has the allele with the 8-bp deletion (VIT_202s0154g00120). Although there was a consensus on the identification of *INP1* as responsible for male sterility, the results for a female sterility gene were not yet unanimous and several genes were proposed as responsible for this trait. One of the most promising genes reported for female sterility was *VviPLATZ* (VIT_202s0154g00070)^10^. As far as we were able to identify, this gene only presented one allele in the hermaphrodite reference genome. In our wild plants this gene was heterozygous in the male and homozygous in the female plants.

The results reported in this paper, provided a comparison between the reference genome (*vinifera*) and *sylvestris*. We have identified SNPs and InDels between the two subspecies and investigated the sex *locus* region, providing a further evidence of which genes may be key players in sex specification of grapevine.

## Discussion

The analysis of the wild-type individuals (*V. v. sylvestris*) reveals a high similarity to PN40024, used as reference genome (*V.v. vinifera*). In fact, 89% of the sequences maps only once to PN40024 genome. 10% of the reads whose pair was lost or map to a different chromosome may be due to the remaining heterozygosity present in the reference genome (93% homozygous^1^), namely, due to the 13 virtual chromosomes created, where heterozygous sequences may reside^1,2^. As referred above, we have identified 4,104,719 heterozygous SNPs and 761,638 heterozygous InDels. A similar study from 2017^19^ comparing four grapevine cultivars obtained a number of heterozygous SNPs ranging between 178,650 and 934,946 and heterozygous InDels of 16,768 – 47,852. The higher number obtained in our study suggests a large distance between wild-type individuals (*sylvestris*) and the domesticated grapevine (*vinifera*). The ratios heterozygous/homozygous are in the same range as calculated by Mercenaro et al. (2017)^19^.

It is described that the domestication pressure set on cultivated varieties, improving the grape characteristics, affected all chromosomes in a similar way^3,20^. In the present study, we were unable to detect a particular chromosome with significantly different number of modifications, measured by a single nucleotide change or Insertion/Deflection. Nevertheless, the chromosomes 6 and 9 were highlighted as the ones with fewer and more alterations, respectively, but the number of modifications are still inside the Tukey’s Fences used on the outlier’s analysis.

As already expected, the majority of the modifications are found on non-coding regions, where the domestication and selection pressures are not as evident as in the coding regions. However, the number of synonymous modifications inside exons is quite similar, the non-synonymous ones being slightly higher. This seems to indicate that the pressures put on *sylvestris* during domestication favoured non-synonymous modifications required for evolution.

One of the most unexpected result is the low Ti/Tv ratio (transitions over transversions) in exons, regardless the SNP impact on protein sequences (synonymous or non-synonymous). Ti/Tv ratios have been associated with the genomic context of the modification and in fact, we clearly observe an association between the genomic context and the ratio. However, the ratio we have obtained on exons (1.4 – 1.9) highly contrasts with the values reported by Wang et al. (2015)^12^ (circa 2.8). A lower ratio as we observed, is getting closer to the ratio that should be observed if SNPs were random (0.5 as there are two transitions and four transversions). Low values of Ti/Tv were also observed in *Oryza sativa*^21^ and *Brassica rapa*^22^. It is evident and intriguing that, in plants, the higher the impact of a SNP, the lower is the Ti/Tv obtained.

Premature start or stop codons, as well as the loss of start and stop codons, may be an indication that these genes are not functional. This genome re-sequencing shows that the number of premature start codons is similar to the number of premature stop codons and the number of start losses is in line with the number of stop losses. This indicates that the overall number of genes effectively relevant in the wild-type is approximately equal to the number of genes present in PN40024: the genes that acquired new start and stop codons, together with the genes with loss of those codons, may represent pseudogenes, differentially relevant for the evolution pattern of each subspecies. Even in InDels, the number of significant alterations is relatively low, what, as suggested by Ramos et al. (2014 and 2017)^14,15^, indicates that both transcriptomes (*sylvestris* and *vinifera*) are mostly similar with only slightly differences.

The fact that the two individuals used in this study share only half of the identified SNPs and InDels, suggests that these individuals are not parentally related. This idea is also supported by the heterozygous over non-reference homozygous SNPs whose ratio has been associated with lineages^12^. We obtained distinct het/nonref-hom ratios for each individual for the same genomic regions. Unfortunately, other *Vitis* studies^11,19^ lacks this kind of analysis.

When considering SNPs and InDels in common (between the two wild individuals), we need to be caution when extrapolating the results to the entire wild population, as we are observing the pattern of only two individuals (one male and one female individuals). Nevertheless, as they were phenotypically distinct individuals (one producing male flowers and the other female flowers), the common modifications may correspond to wild-type specific characteristics.

As both individuals produce distinct flower types, some of the genetic differences found between them may be associated to sex. Curiously, chromosome 2, where the *locus* associated to sex has been reported by different studies^7–9^, has a similar number of SNPs and InDels, when compared to other chromosomes. Nevertheless, we performed a more detailed analysis of the sex-associated *locus* in chromosome 2. From the 46 genes studied on this chromosome, only one (VIT_202s0154g00030) has identical alleles in both wild-type individuals, and other gene was recently associated with male sterility (*INP1*)^10,11^, which makes relevant the question of whether or not the other 44 genes could be associated with sex. At present, we are not able to answer this question and further studies should be carried to determine the impact of these 44 genes, using more individuals of each flower type, in order to better associate each one of these genes with its putative role in flower sex determination.

Considering the relevance of the CRIBI database in *Vitis* research, we need to stress the importance of a complete curation of the genome deposited here. An effort should be carried out to resolve the sequences deposited in the “unknown” and “X_random” chromosomes, as these sequences usually represent alleles of the genes present in the remaining 19 chromosomes. In general, the work described in this article contributes to the identification of putative specific modifications of *sylvestris* when compared to *vinifera*. This paper clarifies questions raised on previous studies using the transcriptome^17,18^, where different levels of gene expression were observed between *sylvestris* and *vinifera* transcriptomes. The identification of SNPs and InDels in the promotor region of the genes (5 kbp upstream of the gene) may explain the differential expression levels detected. In addition, the nucleotide differences revealed here, within coding regions, suggest that different alleles or gene duplications may have distinct importance in *sylvestris* and in *vinifera*.

## Materials and Methods

Sampling and DNA extraction. Representatives from *V. v. sylvestris* (wild type) were selected from INIAV, Dois Portos (Lisbon district, Portugal, 39.041395, −9.181956), where a wild type collection was established. One of the selected individuals exhibited female flowers (WF) and the other displayed male flowers (WM). Fresh leaves were collected from each selected individual. Leaves were grinded in liquid nitrogen and the DNA was extracted and purified with the DNeasy Plant Mini Kit (Qiagen), according to the manufacturer’s instructions. DNA quantity and quality was evaluated using a microplate reader Synergy HT (Biotek, Germany), using the software Gen5™ (Biotek, Germany) and confirmed on a 1.7% (w/v) agarose gel.

### High-throughput sequencing and reference assembly

Genome sequences were obtained individually from each individual using 1 µg of DNA, on a Genome Sequencer Illumina HiSeq 4000 technology (125bp PE; Figure 1). A total of 539,473,738 reads were generated on both sequencings. Low quality reads (phred quality lower than 15) and reads without mate were removed. Additionally, low quality bases were removed from 3’ and 5’ ends. For the WF genome, 216,895,404 reads (91.6% of the total) were used on further analyses, while the WM genome was represented with 277,949,176 high quality reads (91.8%). Considering the *V.v. vinifera* cv. PN40024 reference genome available^1^, we estimated a genome size of about 486 Mbp, for which our predicted coverage ranges between 56X (WF) and 72X (WM). For general proposes, the results obtained from the different individuals were merged together, to better represent a wild-type genome.

The good-quality reads were mapped against the reference genome^1^ using BWA^23^ with the default parameters. About 97% of the input reads mapped correctly to the reference genome (Table 1). Considering the artificial coverage created due to PCR amplification during library preparation, the reads were submitted to Picard (https://broadinstitute.github.io/picard/). If both reads of the same pair mapped against the same genomic location and with the same orientation of other pair, it was considered a duplication and removed. The presence of insertions and deletions, on the new obtained genomes, compared to the reference genome, promoted the occurrence of punctual misalignments. These misalignments were locally realigned with GATK^24,25^. Taking into consideration the reported quality score, sequencing cycle and context, a final Base Quality Recalibration was performed with GATK, to improve the base quality score of the reads. About 92% of the mapped reads were kept after these three refinement steps.

### Variant analyses

SNP and InDel identification between the wild genomes and the reference genome, was performed with GATK’s Haplotype Caller ^24,25^ and annotation according to the gene context was achieved with snpEff (http://snpeff.sourceforge.net), according to PN40024 12X V2.1. Variant quality was estimated with GATK’s Variant Annotator module^24,25^ and false positives were filtered with GATK’s Variant Filtration module^24,25^. Analysis were performed on RStudio (1.0.143)^26^, running over R (3.4.3)^27^. Outliers were estimated based on Tukey’s fences^28^, considering k = 1.5. Venn Diagrams were obtained from calculate.overlap and draw.pairwise.venn functions (VennDiagram package, version 1.6.17^29^).

Transitions over transversion ratio (Ti/Tv) as well as heterozygous over non-reference homozygous SNPs (he/nonref-hom) were calculated for each genome with SNPs grouped by chromosome (discarding the “Un” and “X_random” virtual chromosomes). Calculations were performed with Python (3.7.4) running over Jupyter Notebook (6.0.1). Modules pandas (0.25.1) and matplotlib (3.1.1) were used to manipulate data and draw boxplots.

All scripts may be provided upon request.

### *De novo* assembly

A *de novo* assembly of the genomes was performed in order to obtain the genomic sequences of *loci* that were unclear on the reference genome. From the bulk of sequenced reads (Table 2), an assessment was performed using FastqC software (v0.11.7). Considering the quality indicators, the reads were cleaned with BBDuk (v38.01) from BBMap package. A pipeline was set to remove adapters from both ends (ktrim=r and ktrim=lm; with options k=23, tpe and tbo, using the provided adapters.fa as reference) and low quality reads (options qtrim=rl and trimq=20). After cleaning, the reads quality was re-evaluated on FastqC. Female sample kept 224,542,530 reads while the male sample kept 287,279,296 reads (Table 2). The coverage estimates for this assembly was 58X to female and 74X to male individual (Table 2). Reads were prepared for assembly using fq2fa function from IDBA package^30,31^, with options -- merge and --filter. Assembly was conducted with idba_ud. The contigs obtained with less than 1,000 bp were removed, as this represent low informative segments. The female genome was comprised into 102,650 contigs with a total size of 364,897,899 bp (N50 of 5,795), whereas the male was represented by 108,550 contigs, totalling 370,798,370 bp (N50 of 5,317; Table 3).

**Table 2.** Parameters related to *de novo* assembly of the wild female (WF) and male (WM) individuals. Total number of reads: The number of reads obtained from the sequencing. Reads used on the assembly: Number of reads after cleaning with BBDuk (BBMap package). Coverage: Expected coverage, considering the reference genome of 486 Mbp. Number of contigs: The amount of contigs obtained from the assemblies with more than 1kbp. Total assembly size: The total size (in base pairs) of the obtained genome. N50: The size of the shortest contig at 50% of the total genome length. %CG: Percentage of Cytosine and Guanines over the total genome length.

| ***De novo* assembly parameters** | **WF** | **WM** |
| --- | --- | --- |
| Total number of reads | 236,749,964 | 302,723,774 |
| Reads used on assembly | 224,542,530 | 287,279,296 |
| Coverage | 58X | 74X |
| Number of contigs | 102,650 | 108,550 |
| Total assembly size (bp) | 364,897,899 | 370,798,370 |
| N50 | 5,795 | 5,317 |
| %CG | 34 | 34 |

A %CG ratio of 34 was identified for both samples under study.

Contigs obtained on *de novo* assemblies were queried against the genes annotated on the sex region^7,8^, retrieved from the reference genome (hermaphrodite), on a blastn^32,33^ run with e-value set to 1e-10. The selected contigs were manually inspected. Although the usage of only two individuals, one of each sex, lacks the resolution to differentiate between sex-specific patterns and individual evolutionary mutations, this approach may provide a little insight into the genetic differences leading to different flower types.

The results of this assembly were validated and manually inspected on the region of chromosome 2 that have been associated with sex^7,8^.

### Validation

Four genes involved in flowering development were selected and specific primers were designed (Supplementary Table S1) using Primer Premier 5 (Premier Biosoft). Amplification reactions were performed in 25 µL composed by1 µg of DNA, PCR buffer (20 mM Tris–HCl [pH 8.4], 50 mM KCl), 1.5 mM of MgCl_2_, 0.2 mM of dNTP mix, 0.4 μM of each forward and reverse primers, 5 U of Taq DNA polymerase and autoclaved MiliQ water. The applied program had 4 min for initial denaturation at 94°C; 30 cycles at 94°C 45s (denaturation), annealing at specific temperature (Supplementary Table S1) 45s (annealing) and 72°C 90s (extension); and a final extension at 72°C for 4 minutes. PCR products were purified using PureLink^Tm^ (Invitrogen) and sequenced by Sanger without cloning.

## Data availability

The raw data analysed in this study was submitted to the Sequence Read Archive (SRA) database (www.ncbi.nlm.nih.gov/sra/) under the number PRJNA549602.

## Acknowledgements

We are grateful to Eng. Eiras-Dias, curator of the Portuguese Ampelographic Collection (property of Instituto Nacional de Investigação Agrária e Veterinária, Dois Portos) where sampling was performed, for the collaboration in this work allowing the access to the *Vitis* collection.

We also want to thanks Prof. Helena Oliveira for the support given to genomes re-sequencing. This work was supported by Fundação para a Ciência e Tecnologia (FCT) by granting research centres LEAF (UID/AGR/04129/2013) and BioISI (UID/MULTI/04046/2013). Authors JLCoito, MJNRamos, MRocheta, were funded by FCT fellowships SFRH/BD/85824/2012, SFRH/BD/110274/2015, SFRH/BPD/64905/2009, respectively. MMRCosta was funded by FCT, MCTES and PIDDAC (Portugal) under the project PEst-OE/BIA/UI4046/2014. We received the support of the Research Infrastructure PORBIOTA under the project POCI-01-0145-FEDER-022127.

## Author Contributions

M.J.N.R., J.L.C., D.F.S. and M.R. conceived and designed the experiments. M.J.N.R. and D.F.S. performed and handled bioinformatic analyses. M.J.N.R., D.F.S. and M.R. analysed the data. M.M.R.C. and S.A. critically revised the manuscript. S.A. open the opportunity to develop this work. J.C. established Vitis vinifera sylvestris collection and collected plant tissues. All authors collaborated in the manuscript writing and all reviewed the manuscript.

## Additional Information

### Competing interests

The authors declare no competing interests. Correspondence and requests for materials should be addressed to MJNR or MR.

